# Do pesticide residues have enduring negative effect on macroinvertebrates and vertebrates in fallow rice paddies?

**DOI:** 10.1101/2021.07.06.451252

**Authors:** Jheng-Sin Song, Chi-Chien Kuo

## Abstract

Rice is one of the most important staple food in the world, with irrigated rice paddies largely converted from natural wetlands. The effectiveness of rice fields in help preserve species depends partially on management practices, including the usage of pesticides. However, related studies have focused predominately on the cultivation period, leaving the effects of soil pesticide residues on aquatic invertebrates during the fallow periods little explored; other animals, such as waterbirds, also rely on aquatic invertebrates in flooded fallow fields for their survival. We therefore investigated vertebrates and macroinvertebrates (terrestrial and aquatic) on rice stands and in flooded water during cultivation and fallow periods in organic and conventional rice fields in Taiwan. Association of environmental factors with terrestrial and aquatic organisms was also analyzed. In total, 32,880 individuals of 144 invertebrate families and 381 individuals of 15 vertebrate families were recorded after nine samplings each for six organic and six conventional rice fields. Family richness and abundance of all invertebrates (terrestrial and aquatic) were higher in organic than in conventional fields during the cultivation period, but were similar between the two agricultural practices during the fallow period. Richness and abundance of terrestrial invertebrates in both organic and conventional fields increased with the progression of rice cultivation, so did the differences between the two practices. Richness of aquatic invertebrates was mostly constant across the sampling period, while abundance increased but differences decreased during the fallow period. Richness and abundance of terrestrial invertebrates were positively associated with ambient temperature and height of rice stand. Abundance of aquatic invertebrates were positively associated with pH value and amount of dissolved oxygen but negatively associated with water temperature. Richness and abundance of all vertebrates and each of the constituting groups (fish, amphibian, reptile, bird, and migratory waterbird) were statistically similar between the two practices although abundance of migratory waterbirds in organic fields were two times those in conventional fields during the fallow period. Our study suggested accumulated effects of pesticides on suppressing terrestrial invertebrates during the cultivation period, but diminishing effects of pesticide residues on repressing aquatic invertebrates during the fallow period. This comprehensive study provided a holistic picture on macroinvertebrate and vertebrate fauna, as well as ramifications of pesticide usage, in a representative Southeast Asia rice paddy ecosystem. Further study should compare rice fields with natural wetlands to better assess how to capitalize on agroecosystems for biodiversity conservation.

## 1. Introduction

Agricultural field is one major biome on the planet and farming continues to dramatically transform natural landscapes (Foley et al., 2005). For example, 27% of deforestation over the period 2001-2015 is attributed to commodity production, mainly crop cultivation and cattle grazing (Curtis et al., 2018). To mitigate the detrimental impacts of agricultural intensification on biodiversity and simultaneously raise agricultural productivity (e.g. suppressing agricultural pests or benefiting pollinators), various remedial measures have been proposed, such as reducing pesticide use or diversifying habitats in cultivated and surrounding fields (Gurr et al., 2003; Ricketts, 2004; Bengtsson et al., 2005). For example, the European Union has introduced the agri-environment schemes to safeguard biodiversity and enhance sustainability in farmlands (Batáry et al., 2015).

Rice is one of three most important crops in the world and is the staple food for almost half of the world populations (Prasad et al., 2017). Rice is also the most extensively cultivated crop, grown in over 100 counties of six continents, and covering more than 11% of all arable lands (Donald, 2004; Rao et al, 2017). Among the global rice grown areas, about 90% is located in Asia, especially in East Asia (33%) (Rao et al, 2017). Depending on altitude and source of water, rice field ecosystems can be divided into four categories, including (from high to low elevations) upland, rainfed lowland, irrigated, and flood-prone ecosystems. Irrigated lands account for 75% of global rice production and cover half of global rice fields (Prasad et al., 2017). These irrigated lands are mostly converted from wetlands, leading to a great loss in natural wetlands (Donald, 2004). However, these flooded rice paddies can potentially be surrogate habitats for wetland species, such as waterbirds, and can help mitigate the negative influence of habitat loss (Fasola and Ruiz, 1996; Lawler, 2001; Czech and Parsons, 2002; Elphick and Oring, 2003; Toral and Figuerola, 2010; Herring et al., 2019; Kasahara et al., 2020). The effectiveness of rice fields in sustaining species depends partially on management practices (Tourenq et al., 2003; Elphick et al. 2010; Strum et al., 2013), including the usage of pesticides (Simpson and Roger, 1995; Parsons et al., 2010). For example, effects of pesticide use on aquatic invertebrates in rice paddies generally found decreased species diversity but increased abundance of primary consumers (Simpson and Roger, 1995; Suhling et al., 2000; Wilson et al., 2008; Kumar et al., 2013; Stenert et al., 2018). However, these studies were predominately implemented during the cultivation periods. It is unclear whether pesticide residues in soils (Gevao et al., 2000) can have any lasting effect on aquatic invertebrates during the fallow periods when pesticides are no longer applied. This is important especially when other animals, such as waterbirds, also rely on aquatic invertebrates in flooded fallow fields for their survival (e.g. Fujioka et al., 2001, 2010; Stafford et al. 2010; Katayama et al., 2020).

In Taiwan, rice is the most important crop and occupies half of arable land. The majority of fields is cultivated with the japonica variety (87%, *Oryza sativa* subsp. *japonica*), followed by the indica variety (8%, *O. sativa* subsp. *indica*) (Hsing, 2016), and is predominately planted in flooded instead of dry fields. Given the well documented harmful effects of pesticides on farmland animals, organic farming practice free of pesticide usage is advocated against conventional farming practice that keeps applying pesticides. This is followed with numerous studies comparing a diverse set of animal taxa between organic and conventional farmlands (e.g. Mäder et al., 2002; Bengtsson et al., 2005; Hole et al., 2005; Rizo-Patrón et al., 2013; Reganold and Wachter, 2016; Toffoli and Rughetti, 2017; Katayama et al., 2019). This is no exception for Taiwan that focuses particularly on rice paddy ecosystems. Fan (2016) and Sun (2020) compared arthropods on rice stands in organic vs. conventional paddies in eastern and western Taiwan, respectively. Fan (2016) found higher species richness and abundance in organic fields, but Sun (2020) reported no difference between the two practices. There was no difference in species richness of four functional groups but higher abundance of predators in organic paddies for arthropods on rice stands in western Taiwan (Huang et al., 2020). Also in western Taiwan, biomass proportion of three arthropod functional groups on rice stands and in flooded water differed between organic and conventional paddies during the cultivation period (Huang et al., 2018b). Lastly, species richness and abundance of soil nematodes were similar between organic and conventional rice paddies in western Taiwan (Chen, 2018). These studies targeted predominately on arthropods on rice stands during the cultivation period and focused mainly on ecosystem services provided by predators and parasitoids in suppressing agricultural pests.

In northern Taiwan where the fall temperature is not high enough for completing another round of rice cultivation, the rice field is left fallow from August to February but is still flooded (depth 10-30 cm) to control weeds. This is different from other parts of Taiwan where rice is typically cultivated twice a year, from February to June and from July to November, respectively. The uncultivated but flooded rice fields in northern Taiwan can provide supplementary habitats for wetland species, such as migratory birds that arrive in Taiwan from October to March (Lai, 2012; Lu, 2019). In this study, we compared animal taxa between organic and conventional rice paddies during both cultivation and fallow periods. Both rice stands and flooded water were sampled for terrestrial and aquatic organisms. Animals investigated includes not only arthropods but also other macroinvertebrates (e.g. mollusks, annelids, etc.) and vertebrates. In addition, association of richness and abundance of invertebrates with environmental factors was examined. We hypothesized that (1) species richness and abundance will be higher in organic than in conventional paddies; (2) the differences in species richness and abundance between the two agricultural practices will increase with the progression of rice cultivation (after continual pesticide application in conventional fields) but decrease following the onset of fallow period (after the termination of pesticide usage in conventional fields); (3) the number of migratory waterbirds (family Anatidae, Charadriidae, and Scolopacidae) will be similar between the two practices as they arrived in the middle to late stage of fallow period when pesticide residues in the water were largely diluted. Despite that rice is primarily cultivated in Asia, the benefits of organic rice farming to biodiversity are largely studied in Europe and North America (Amano, 2009; Katayama et al., 2019). Additionally, most previous studies focused either on vertebrates or macroinvertebrates, sometimes further divided into terrestrial or aquatic invertebrates, and considered only cultivated or fallow period. In comparison, our comprehensive study should provide a more holistic picture on ramifications of pesticide usage to rice ecosystems in Asia.

## 2. Materials and methods

### 2.1. Study area

The study was conducted from March 2019 to March 2020 in Yuanshan and Sanxing of Yilan county in northeastern Taiwan (Fig. 1). Yilan is one of the main rice cultivation area in northern Taiwan (Agricultural Statistics Yearbook 2019). A total of 12 sites, including six organic and six conventional rice paddies, were surveyed for macroinvertebrates and vertebrates. However, because three of the organic fields were found devoid of water during the fallow period (these three sites were nevertheless flooded for the previous fallow period, so they were initially selected for survey), another three new organic rice fields of similar area (*t* = 0.9, *P* > 0.05) were instead sampled to the end of the study (Fig. 1). Sites were surrounded by farmlands of the same agricultural practices (i.e. organic or conventional) and at least two of the four surrounding banks were made of soil instead of concrete. In addition, sites were away from human settlements and main roads and distances between sites were >100 meters. Organic rice fields were located by referring to the geographic information system maintained by the Yilan county (https://ilanland.e-land.gov.tw/) and were confirmed by inquiring local farmers. There was no significant difference in area between organic and conventional fields (averaged 0.24±0.10 and 0.20±0.07 (±SD) ha, respectively; *t* = 0.9, *P* > 0.05). In Yilan, pesticides are commonly applied to conventional rice paddies thrice: in late February (shortly before the onset of cultivation), March-April, and May-June (Huang, 2008).

**Fig. 1.**
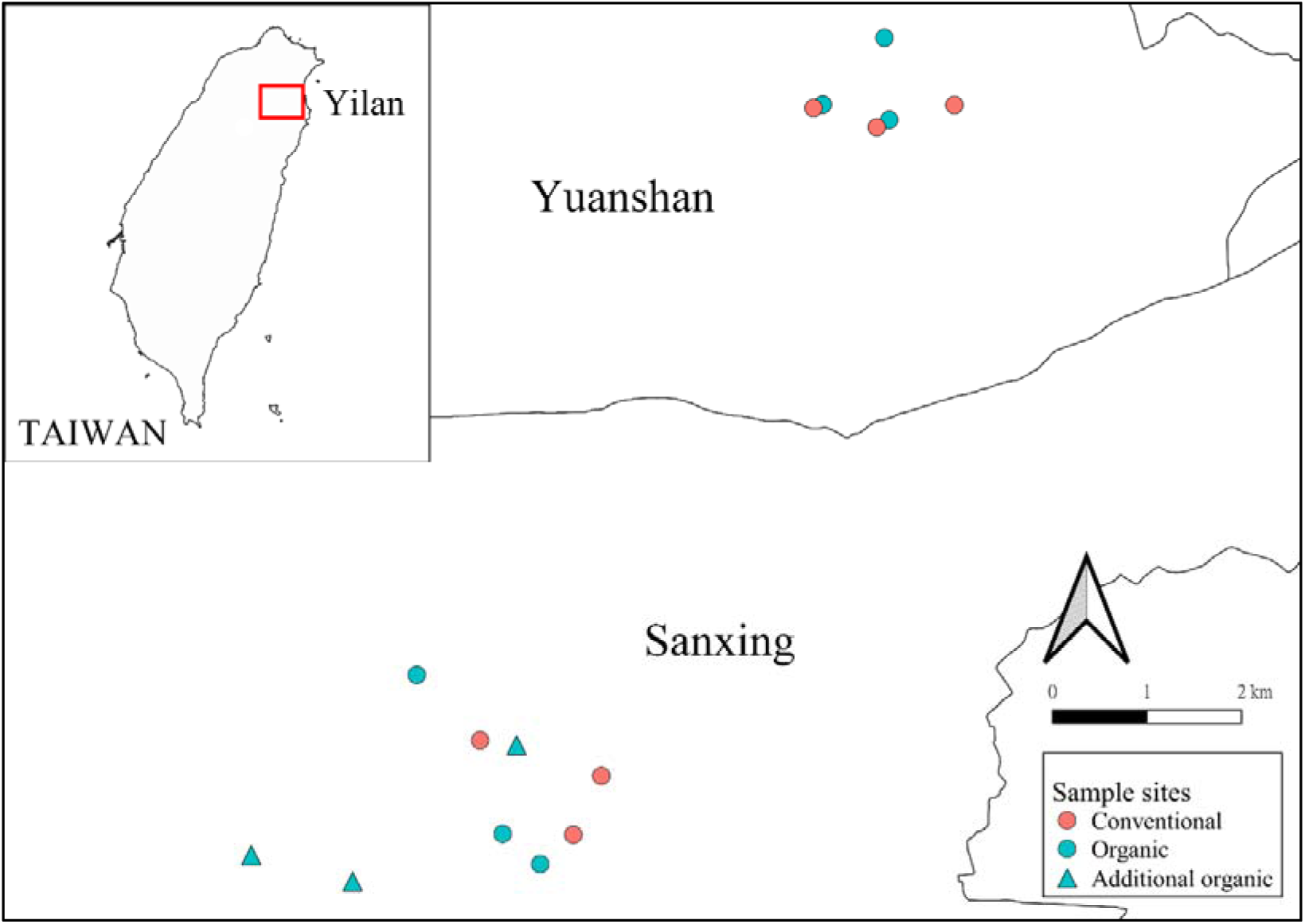
Study area in Yilan County, Taiwan.

### 2.2. Animal sample collection

Each site was sampled for a total of nine times, including five during the cultivation period and four during the fallow period (Fig. 2). Organic and conventional fields were surveyed alternatively during each sampling session, which was typically finished within three days to control for potential temporal variation (inclement weather has once protracted the session for two weeks, twice for one week). The number and identity of vertebrates and large flying arthropods (e.g. dragonfly and butterfly) were recorded upon visiting the site (except for fish). Birds were surveyed across the whole site by using a binocular (10×32, Optisan BRITEC CR). Because birds were not always surveyed at dawn or dusk when they are typically more active, we may underestimate bird abundance. However, the goal to compare difference between the two agricultural practices can still be fulfilled with no bias in our surveillance time (e.g. without one always surveyed at dawn and the other at noon). Amphibians, reptiles, and large flying arthropods were recorded directly by walking slowly along the full length of the four banks, each enclosing fifty-centimeter belts extending into the paddy. For macroinvertebrates and fish on rice stands and in flooded water, we set up four quadrat plots, each with 1 square meter, along the middle of the four banks. Terrestrial and aquatic invertebrates (and fish) were separately surveyed using a nylon sweep net of 36 cm in diameter and a sieve size of 0.25 mm. Each plot was swept four times in reverse 8 shape (separately on rice stands and in flooded water). For some sampling visits, only terrestrial or aquatic animals were collected, depending on whether rice stands existed (cultivation vs. fallow periods) and if the fields were flooded (Fig. 2).

**Fig. 2.**
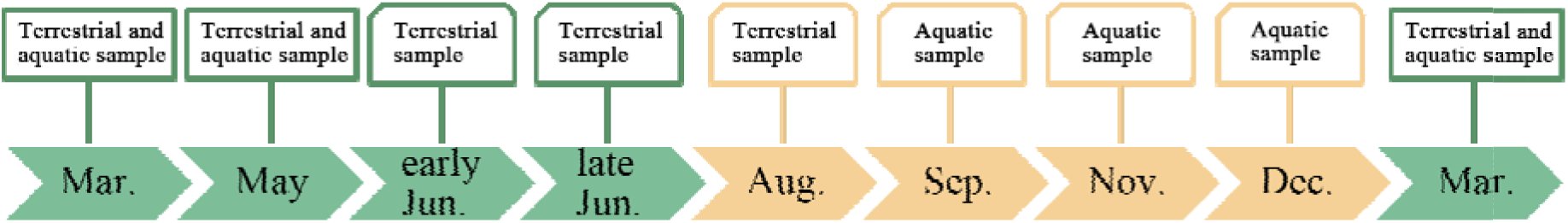
Schedule for sampling terrestrial and aquatic animals in rice fields during the study period from March 2019 to March 2020. Green and orange blocks represented samplings during the cultivation and fallow periods, respectively.

Vertebrates, large flying arthropods, and large gastropods such as Ampullariidae and Viviparidae were identified and counted in the field. The other macroinvertebrates were preserved in 95% ethanol for latter identification in the laboratory. Both vertebrates and invertebrates were identified at least to the family level following published keys (Triplehorn et al., 2005; Chen, 2011; Ferris, 2011; Luo, 2017; Lai and Chen, 2018; Shultz, 2018). Functional groups each organism belonged to were also classified according to food habits and ecological functions (Cummins et al., 2005; Triplehorn et al., 2005; Gerlach et al., 2013). In this study, functional groups included bacterivore, collector, detritivore, fungivore, herbivore, omnivore, parasitoid, pest, pollinator, predator, scavenger, and visitor (Appendix A).

### 2.3. Environmental factors

We took 200 ml water sample each from the same quadrat plot as surveying macroinvertebrate to measure water quality (AZ water quality meter, 86031, AZ Instrument Corp., Taichung, Taiwan). Each plot (4 plots in each site) was measured separately. Water quality included water temperature (°C), pH value, conductivity (μS/cm) and amount of dissolved oxygen (mg/L). The depth of water (cm) and height of rice stand (cm) were measured with a tape. Ambient temperature (°C) of each site was also recorded during each visit.

We also classified banks into four groups according to height and status of the vegetation: 1. no vegetative cover or very sparse; 2. average height <30 cm; 3. average height >30 cm and not wilted; 4. average height >30 cm and wilted.

### 2.4. Data analysis

Site was the independent unit for statistical analyses. Biotic data (e.g. richness) from the four sampling belts or plots were aggregated to the site level data. On the other hand, abiotic data (e.g. water temperature) was averaged across the four plots to attain a mean site value.

Difference between organic and conventional paddies in richness, abundance, Shannon index (all three in family level), and functional groups was assessed with *t*-test or Wilcoxon rank-sum test depending on whether assumption of normality and homoscedasticity were fulfilledt. Effect size between the two agricultural practices (organic relative to conventional) was calculated with Cohen’s *d*, so that when *d* > 0, the value of organic is larger than that of conventional and vice versa. The magnitude of difference is large when the absolute value of *d* is > 0.8, middle when *d* > 0.5, and small when *d* > 0.2.

Difference in family composition of terrestrial and aquatic invertebrates between agricultural practices was each investigated with non-metric multidimensional scaling (NMDS), followed by permutational multivariate analysis of variance (PERMANOVA). Difference in proportions of functional groups was assessed with Fisher-Freeman-Halton’s test with 100,000 Monte Carlo permutations. Association of environmental factors with richness and abundance of invertebrates (separately for terrestrial and aquatic organisms) was examined with generalized estimating equations (GEE) with Poisson distribution and log-link function, followed by Tukey HSD post hoc test. Different samplings in the same site were regarded as repeated measures. All the analyses were conducted in vegan, iNEXT, lme4, geepack, emmeans packages of R (4.0.0).

## 3. Results

Vertebrate and large flying invertebrates were sampled for nine times, whereas terrestrial (on rice stands) and aquatic (in flooded water) macroinvertebrate was each sampled for six times. Terrestrial sampling included five times during the cultivation period and one time during the fallow period. Aquatic sampling included three times each during the cultivation and fallow period (Fig. 2). However, aquatic samples of four sites, two for each practice, could not be collected due to insufficient water levels in May 2019.

### 3.1. Macroinvertebrate family richness, abundance, and diversity

In total, 32,880 individuals of 144 invertebrate families were recorded (Appendix B). We recorded 7,269 terrestrial individuals of 106 families and 11,939 aquatic individuals of 45 families in organic fields; on the other hand, we recorded 4,820 terrestrial individuals of 95 families and 8,852 aquatic individuals of 43 families from conventional fields.

Overall, family richness and abundance of macroinvertebrates were significantly higher in organic than in conventional fields (*t* = 3.3, 3.6, respectively; both *p* < 0.01), but there was no difference in Shannon index between the two practices (*p* > 0.05) (Fig. 3a-c). Separately, terrestrial invertebrate richness and aquatic invertebrate abundance were significantly higher in organic than in conventional fields (*t* = 2.8, *W*= 31, respectively; both *p* < 0.05). Although aquatic richness and terrestrial abundance were similar between the practices (both *p* > 0.05), the effect sizes (organic relative to conventional) were large (*d* = 0.8, 1.3, respectively). On the other hand, terrestrial and aquatic Shannon indices were similar between the two practices (both *p* > 0.05), with small and medium effect sizes (*d* = 0.41, −0.66) (Fig. 3a-c).

**Fig. 3.**
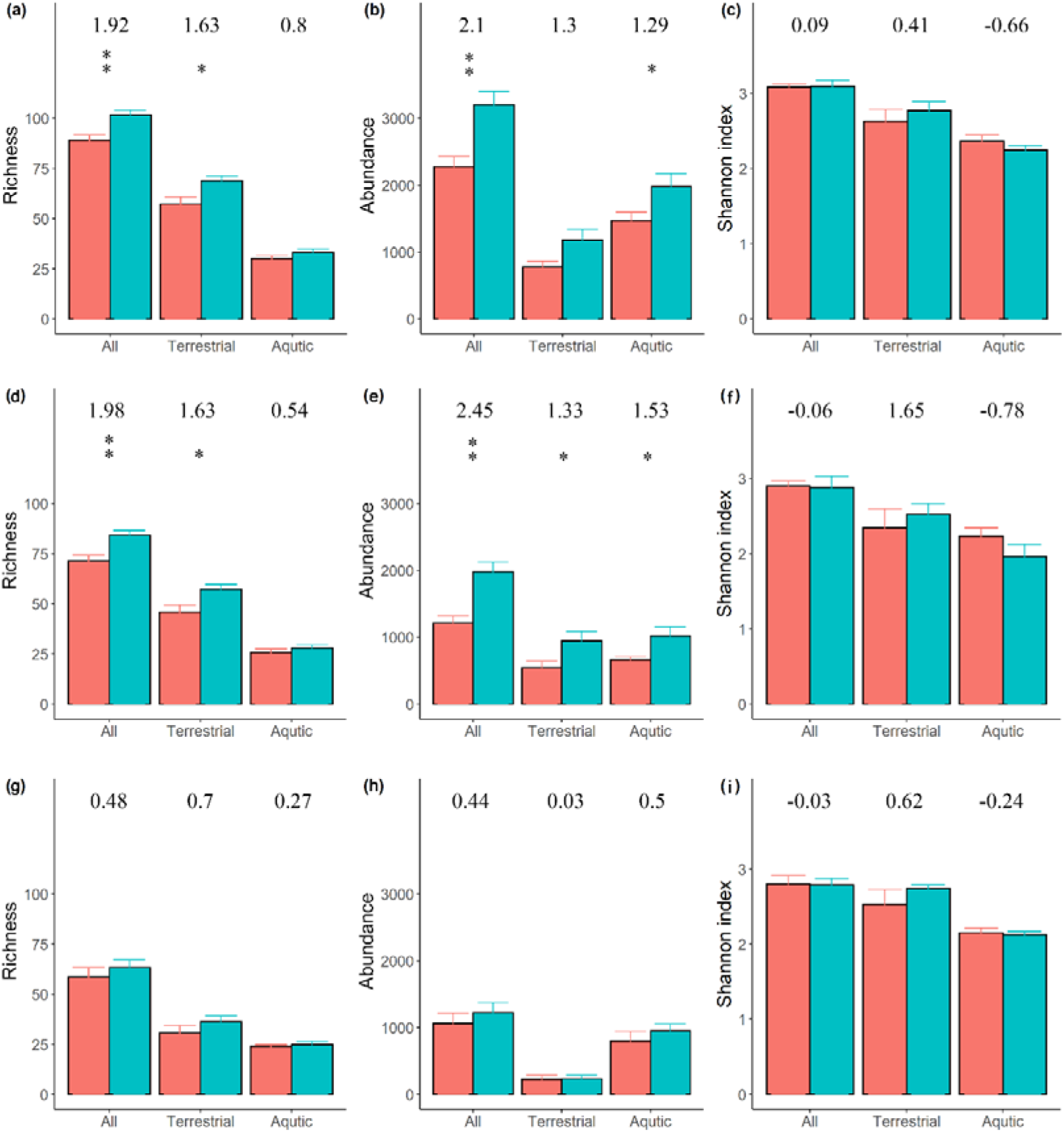
Family richness, abundance, and Shannon index of terrestrial and aquatic macroinvertebrates (mean + 1 SE) in conventional (red) vs. organic (blue) rice fields. (a)-(c) whole period (cultivation + fallow); (d)-(f) cultivation period; (g)-(i) fallow period. \**P* < 0.05, \*\**P* < 0.01. Value above each bar represented the effect size (organic relative to conventional).

We further divided the samples into cultivation and fallow periods. During the cultivation period, both richness and abundance of all invertebrates were significantly higher in organic than in conventional fields (*t* = 3.4, 4.3, respectively; both *p* < 0.01). The richness and abundance of terrestrial invertebrates were also significantly higher in organic fields (*t* = 2.8, 2.3, respectively; both *p* < 0.05). The abundance of aquatic invertebrate was significantly higher in organic fields (*t* = 2.65; *p* < 0.05), but there was no difference for the richness (*t* = 0.9; *p* > 0.05). Shannon indices were similar between the practices for terrestrial, aquatic, and overall invertebrates (all *p* > 0.05), although the effect sizes were opposite for terrestrial and aquatic invertebrates (1.65, −0.78, respectively) (Fig. 3d-f). On the other hand, during the fallow period, richness, abundance, and Shannon index were similar between the organic and conventional fields for terrestrial, aquatic, and overall invertebrates (all *p* > 0.05) although the effect sizes were largely positive except for Shannon indices for aquatic and overall invertebrates (Fig. 3g-i).

Regarding temporal variation for terrestrial invertebrates, richness and abundance in both organic and conventional fields increased with the progression of rice cultivation from March to late June (Fig. 4a-b). Differences in abundance and richness between the two practices also increased with time, from no difference initially to significantly higher in organic fields in late June (*W* = 31, 33, respectively; both *p* < 0.05). The differences then declined during the fallow period in August and the beginning of another cultivation round in March of the following year (Fig. 4a-b). On the other hand, there were no differences in richness and abundance of aquatic invertebrates between the two practices during the cultivation and fallow periods (all *p* > 0.05) (Fig. 4c-d). The richness of aquatic invertebrates was mostly constant across the sampling periods. The effect size increased during both the cultivation (from −0.24 to 0.94) and fallow (from −0.18 to 0.42) period (Fig. 4c). The abundance of aquatic invertebrates decreased with advancement of rice cultivation, but increased during the fallow period. The effect size increased during the cultivation period (from 0.74 to 1.32) but decreased during the fallow period (from 0.84 to 0.1) (Fig. 4d).

**Fig. 4.**
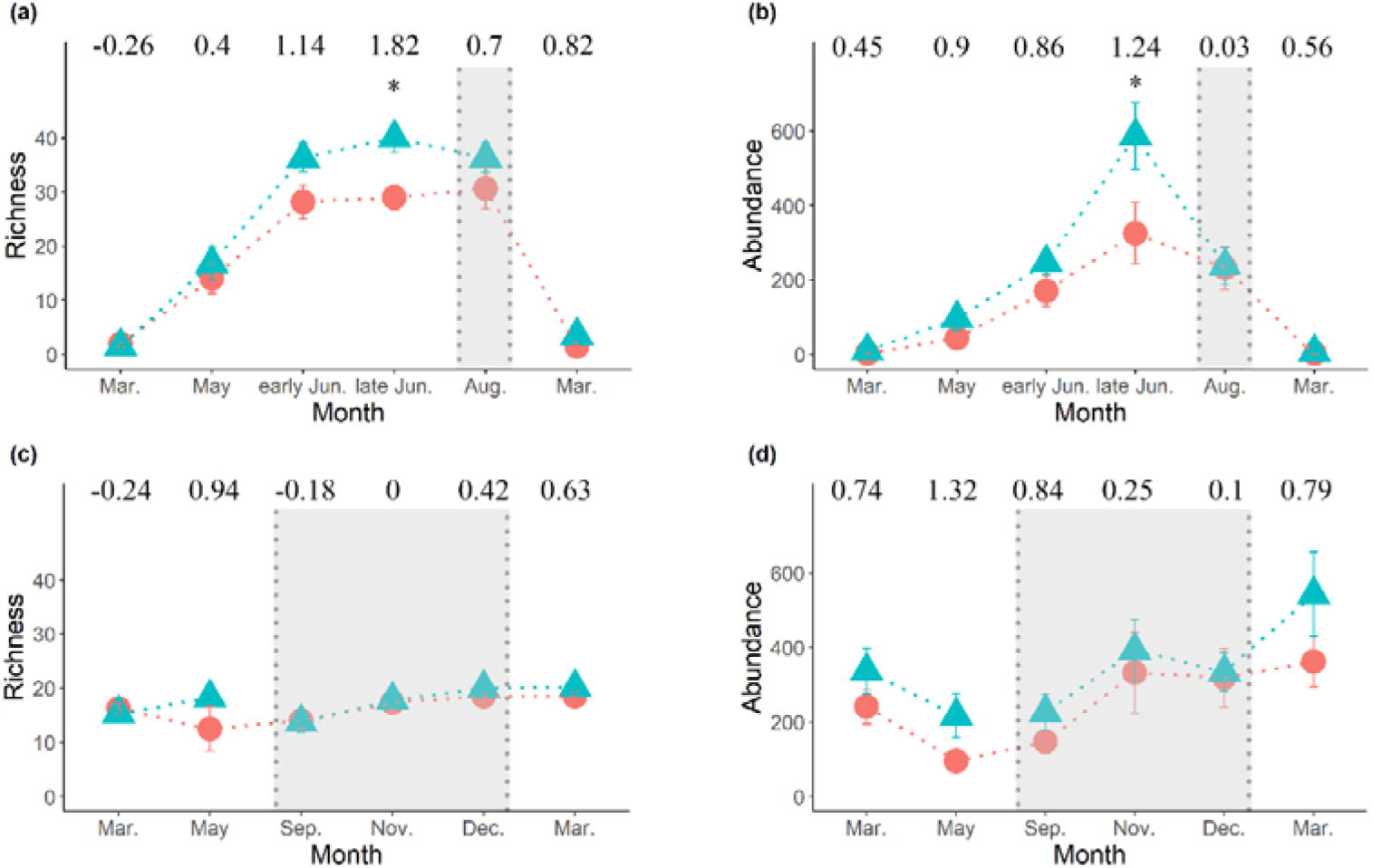
Temporal variation in family richness and abundance (mean ± 1 SE) of terrestrial and aquatic macroinvertebrates in conventional (red) and organic (blue) rice fields from March 2019 to March 2020. (a) terrestrial richness; (b) terrestrial abundance; (c) aquatic richness; (d) aquatic abundance. Shaded areas represented fallow period. \**P* < 0.05. Value above each sample represented the effect size (organic relative to conventional).

### 3.2. Macroinvertebrate family composition

Terrestrial invertebrates were dominated by Delphacidae (22.8%), Chironomidae (17.5%), Cicadellidae (12.8%), Phoridae (4.3%), and Ephydridae (4.2%) in organic fields (totaling 61.6%), while were dominated by Delphacidae (33.0%), Chironomidae (14.3%), Cicadellidae (8.2%), Aphididae (4.7%), and Mycetophilidae (4.4%) in conventional fields (64.6%). There was no significant difference in composition between the two farming practices (PERMANOVA, *p* > 0.05) (Fig. 5a), including separating into cultivation or fallow period (both *p* > 0.05) (Fig. 5c,e).

**Fig. 5.**
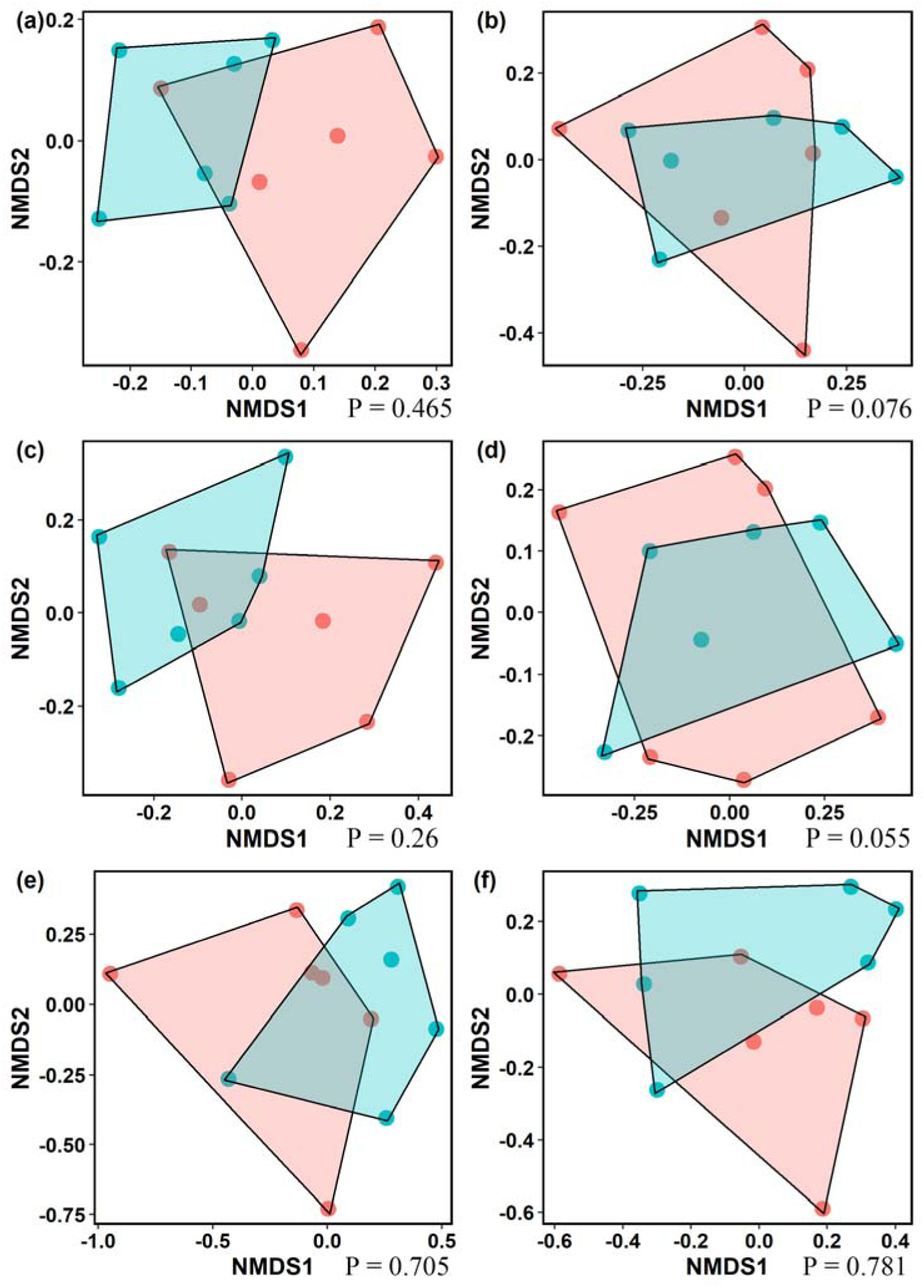
Plots of non-metric multidimensional scaling (NMDS) and *p*-value after permutational multivariate analysis of variance for terrestrial and aquatic macroinvertebrate family composition in conventional (red) vs. organic (blue) rice fields. (a) terrestrial macroinvertebrates in whole period (cultivation + fallow); (b) aquatic macroinvertebrates in whole period; (c) terrestrial macroinvertebrates in cultivation period; (d) aquatic macroinvertebrates in cultivation period; (e) terrestrial macroinvertebrates in fallow period; (f) aquatic macroinvertebrates in fallow period.

On the other hand, aquatic invertebrates were dominated by Chironomidae (26.7%), Micronectidae (14.3%), Naididae (11.4%), Ampullariidae (8.6%), and Dytiscidae (7.8%) in organic fields (totaling 68.9%), while were dominated by Micronectidae (21.0%), Chironomidae (14.0%), Naididae (10.0%), Ampullariidae (8.7%), and Dytiscidae (8.0%) in conventional fields (61.6%). Likewise, there was no significant difference in composition between the two farming practices (*p* > 0.05) (Fig. 5b), including cultivation or fallow period, separately (both *p* > 0.05) (Fig. 5d,f).

### 3.3. Macroinvertebrate functional groups

Organic fields were comprised of collectors (26.7%), pest (21.4%), predators (14.3%), omnivores (10.0%), herbivores (9.3%), scavengers (8.1%), pollinators (7.2%), parasitoids (2.7%) and others (0.3%), while conventional fields were comprised of pest (23.0%), collectors (19.7%), omnivores (15.3%), herbivores (14.6%), predators (13.4%), scavengers (6.0%), pollinators (5.7%), parasitoids (1.8%) and others (0.5%). There was no significant difference in proportions of functional groups between the two practices (Fisher-Freeman-Halton’s test, *p* > 0.05). However, family richness of herbivores and parasitoids was significantly higher in organic than in conventional fields (*W* = 31, 30, respectively; both *p* < 0.05), with high effect sizes (1.49, 1.14). Although richness of predators and pest was not different between the practices (*W* = 30, 30, respectively; both *p* > 0.05), both had large effect sizes (1.1, 0.98) (Table 1). On the other hand, abundance of parasitoids and collectors was significantly higher in organic than in conventional fields (*t* = 2.7, 2.4, respectively; both *p* < 0.05) and both with high effect sizes (1.57, 1.39). Despite that abundance of predators was not different between the practices (*t* = 1.7, *p* > 0.05), the effect size was large (0.99) (Table 1).

**Table 1.** Mean family richness and abundance of different macroinvertebrate functional groups in organic and conventional rice fields, as well as the difference. Effect size (organic relative to conventional; >0.2: small; >0.5: medium; >0.8: large).

| Functional Group | Richness (family) |  | W | Effect size | Abundance |  | *t* | Effect size |
| --- | --- | --- | --- | --- | --- | --- | --- | --- |
|  | Organic | Conventional |  |  | Organic | Conventional |  |  |
| Collector | 10.5 | 9.7 | 26.5 | 0.8 | 853.3 | 447.7 | 2.4* | 1.39 |
| Detritivore | 1 | 1 | 18.0 | 0 | 2 | 3.33 | -0.7 | -0.38 |
| Herbivore | 15.7 | 12.8 | 31.0* | 1.49 | 297.3 | 349.7 | -0.5 | -0.31 |
| Omnivore | 6.6 | 6.3 | 20.5 | 0.17 | 318.7 | 332.2 | -0.1 | -0.05 |
| Parasitoid | 11.3 | 9.2 | 30.5* | 1.14 | 87 | 41.3 | 2.7* | 1.57 |
| Pest | 15 | 13 | 29.5 | 0.98 | 684.5 | 523.7 | 1.4 | 0.79 |
| Pollinator | 4.7 | 4 | 22.0 | 0.5 | 230.8 | 136.8 | 1 | 0.58 |
| Predator | 26 | 23.2 | 30.0 | 1.1 | 458.3 | 306 | 1.7 | 0.99 |
| Scavenger | 8.3 | 7.2 | 25.5 | 0.78 | 260 | 130.3 | 1 | 0.6 |
| Visitor | 1.5 | 1.7 | 13.5 | -0.24 | 7.5 | 6.7 | 0.3 | 0.79 |
| Bacterivore | 0.2 | 0.3 | 15.0 | -0.36 | 0.3 | 0.3 | 0 | 0 |
| Fungivore | 0.8 | 0.5 | 22.5 | 0.5 | 1.5 | 0.7 | 1.1 | 0.64 |

\* $P < 0.05$

In organic fields, richness of terrestrial predators and parasitoids increased throughout the cultivation period, but that of pest and pollinators decreased in the final stage (Fig. 6a). In conventional fields, richness of parasitoids rose with the cultivation, while that of predators, pest, and pollinators dropped in the end of the cultivation period (Fig. 6b). Abundance of terrestrial functional groups generally increased with the progression of the cultivation period in organic fields (Fig. 6c). The pattern is less consistent in conventional fields, with steady increase in pest and parasitoids throughout the cultivation period but a decline in predators and pollinators during the final stage (Fig. 6d).

**Fig. 6.**
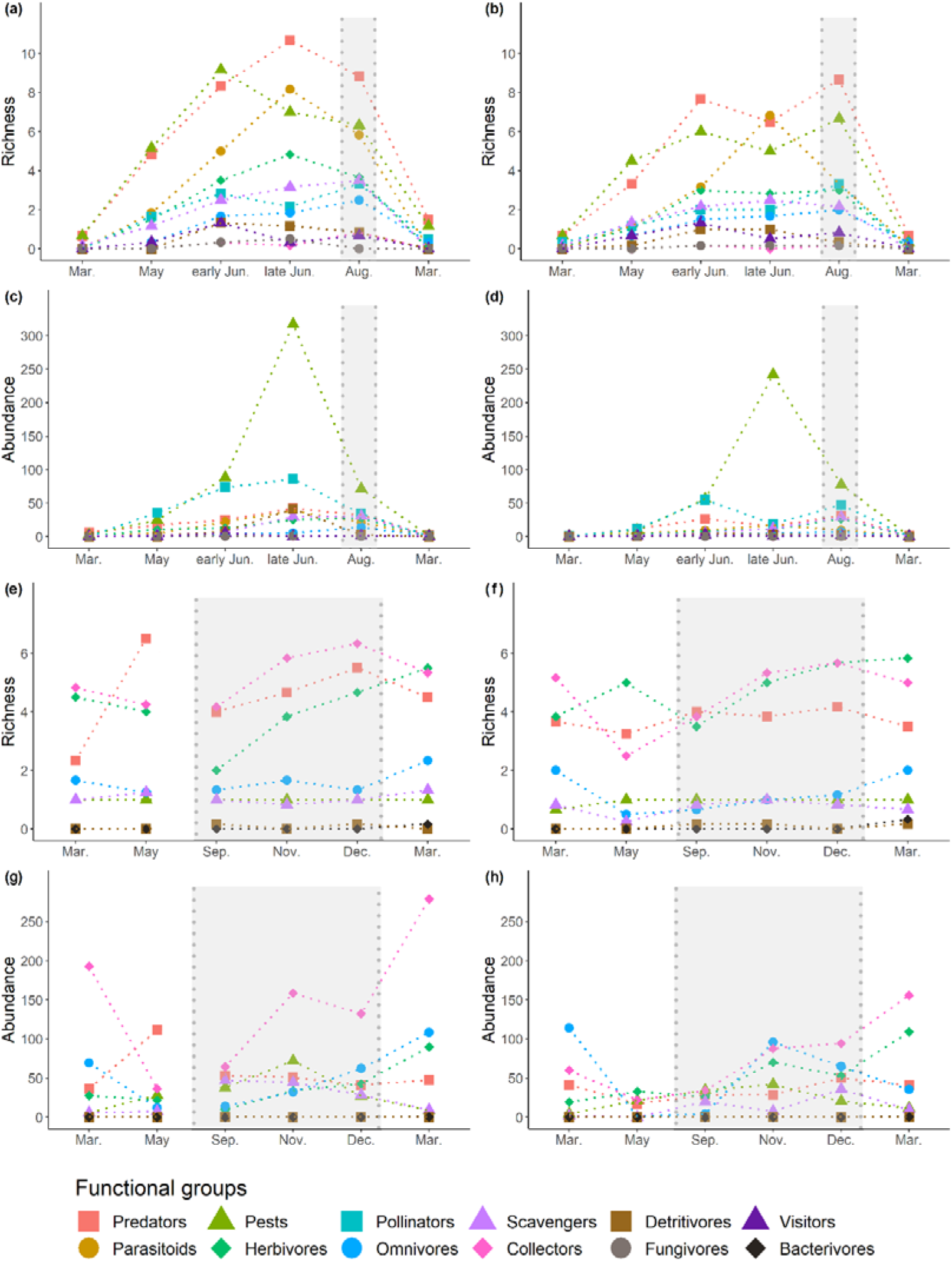
Temporal variation in family richness and abundance (mean value) of different functional groups for terrestrial and aquatic macroinvertebrates in conventional and organic rice fields from March 2019 to March 2020. (a) terrestrial richness in organic; (b) terrestrial richness in conventional;(c) terrestrial abundance in organic; (d) terrestrial abundance in conventional; (e) aquatic richness in organic; (f) aquatic richness in conventional; (g) aquatic abundance in organic; (h) aquatic abundance in conventional. Shaded areas represented fallow period.

On the other hand, temporal changes in abundance and richness of aquatic functional groups in both farming practices were less consistent (Fig. 6e-h). Richness of aquatic predators increased while that of pest largely maintained constant during both cultivation and fallow periods in organic fields (Fig. 6e). Richness of predators and pest kept principally constant over both periods in conventional fields (Fig. 6f). During the cultivation period, abundance of predators decreased and that of pest increased in both field types, (Fig. 6g,h). During the fallow period, abundance of predators and pest increased first, followed by a decrease in organic fields (Fig. 6g). In conventional fields, the trend was similar for pest, but there was a consistent increase in predators over the fallow period (Fig. 6h).

### 3.4. Association of environmental variables with macroinvertebrate family richness and abundance

A total of 108 data (12 sites x 9 surveys) each for ambient temperature, height of rice stand, and bank vegetation type were recorded. Proportions of the four bank types (no vegetation, height < 30 cm, height > 30 cm and not wilted, height > 30 cm and wilted) in organic and conventional fields were 3.7%, 48.2%, 48.2%, 0%, and 27.8%, 18.5%, 18.5%, 35.2%, respectively. Richness of terrestrial invertebrates was affected by all factors except for township (Table 2). Ambient temperature and height of rice stand were positively associated with richness (Fig. 7a,b). Richness was higher in organic than in conventional fields (69.0±2.2, 57.3±3.5, respectively), and higher when bank vegetation was > 30 cm than when it was < 30 cm (Fig. 7c). Abundance of terrestrial invertebrates was positively associated with ambient temperature and height of rice stand (χ^2^ = 65.7, 13.7, respectively; both *p* < 0.001) (Fig. 7d,e), but not agricultural practices, bank vegetation type, and township (Yuanshan or Sanxing) (all *p* > 0.05, Table 2).

**Table 2.**
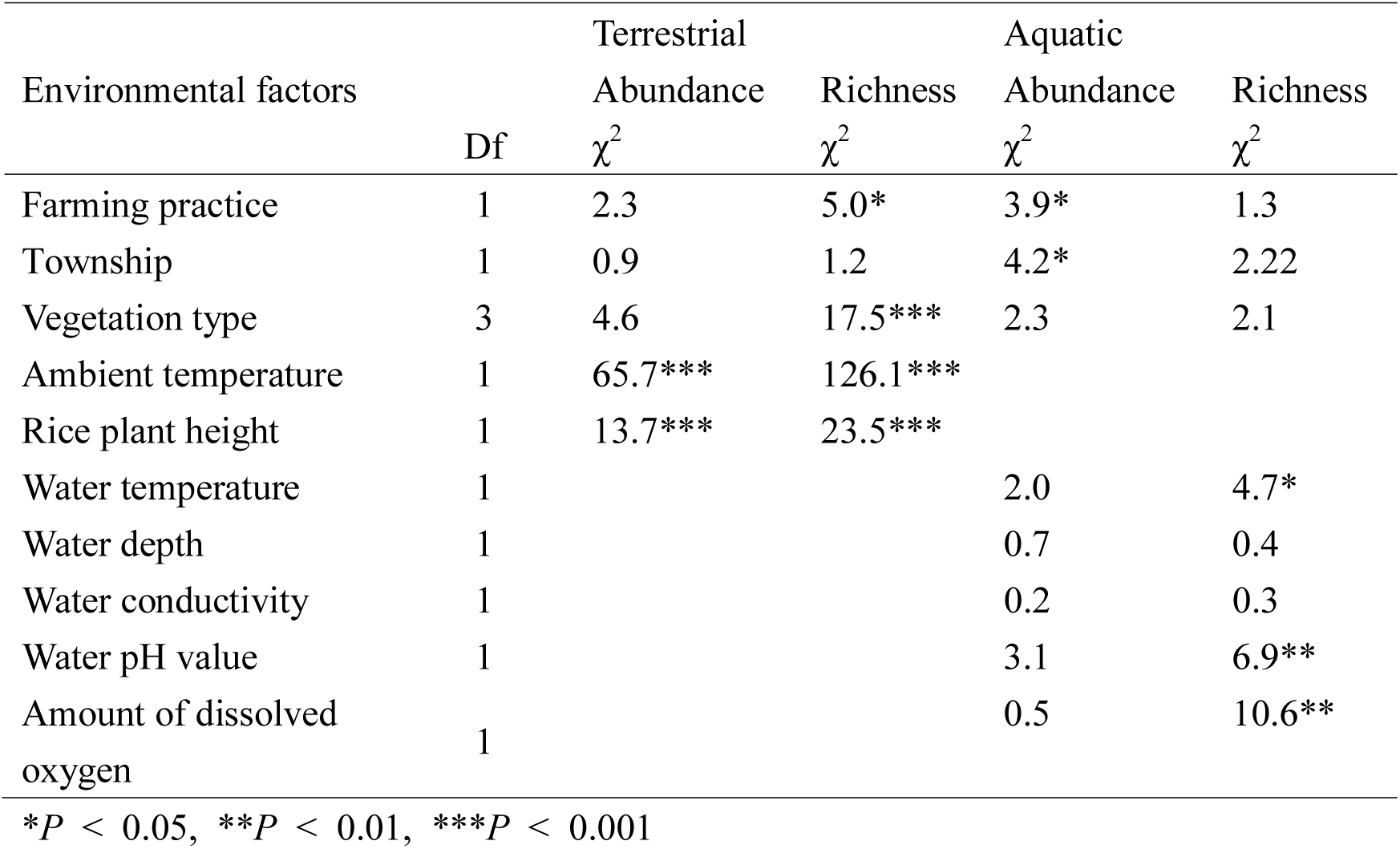
Statistical results for effects of environmental factors on abundance and richness of terrestrial and aquatic macroinvertebartes.

**Fig. 7.**
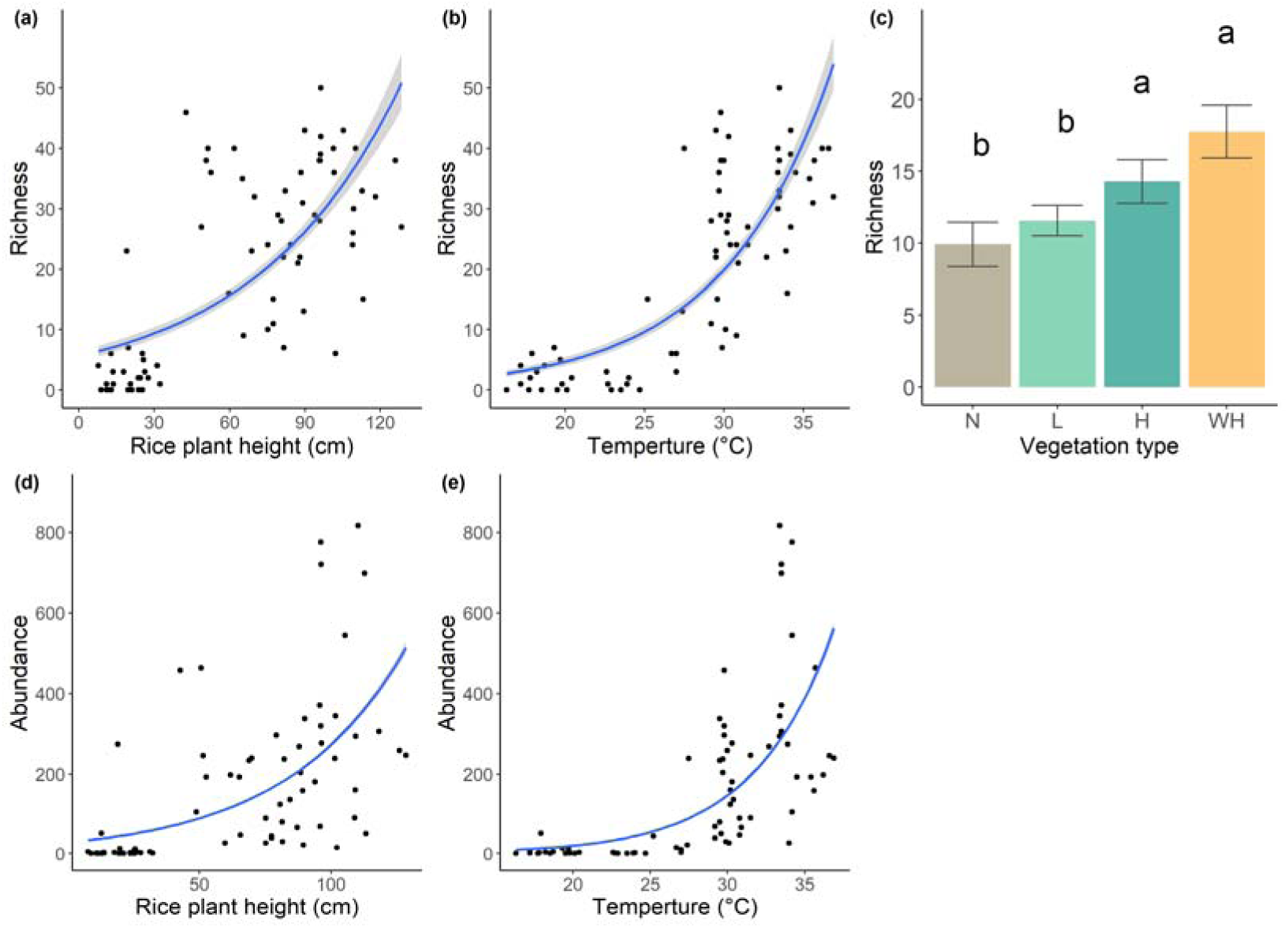
Statistically significant (*P* < 0.05) effects of environmental factors on family richness and abundance of terrestrial macroinvertebrates. Richness: (a) rice plant height; (b) ambient temperature; (c) bank vegetation type. Abundance: (d) rice plant height; (e) ambient temperature. Different letters in (c) represented significant difference.

A total of 272 records each for water depth, water temperature, pH value, conductivity, and amount of dissolved oxygen were documented. Except for dissolved oxygen that was higher in conventional than in organic fields (*t* = −2.25, *p* < 0.05; 4.6±0.3, 5.4±0.3, respectively), values of the other four variables were similar between the farming practices (all *p* > 0.05). Richness of aquatic invertebrates was affected by water temperature, pH value, and amount of dissolved oxygen (χ^2^= 4.7, 6.9, 10.6, respectively; *p* < 0.05, < 0.01, < 0.01), but not the other factors (all *p* > 0.05) (Table 2). Richness was positively associated with pH value and amount of dissolved oxygen, and negatively associated with water temperature (Fig. 8a-c). Abundance of aquatic invertebrates was affected by agricultural practice and township (χ^2^ = 3.9, 4.2, respectively; both *p* < 0.05), but not bank vegetation and the five water characteristics (all *p* > 0.05) (Table 2). Abundance was higher in organic than in conventional fields (1978.7±189.4, 1466.7±130.6, respectively).

**Fig. 8.**
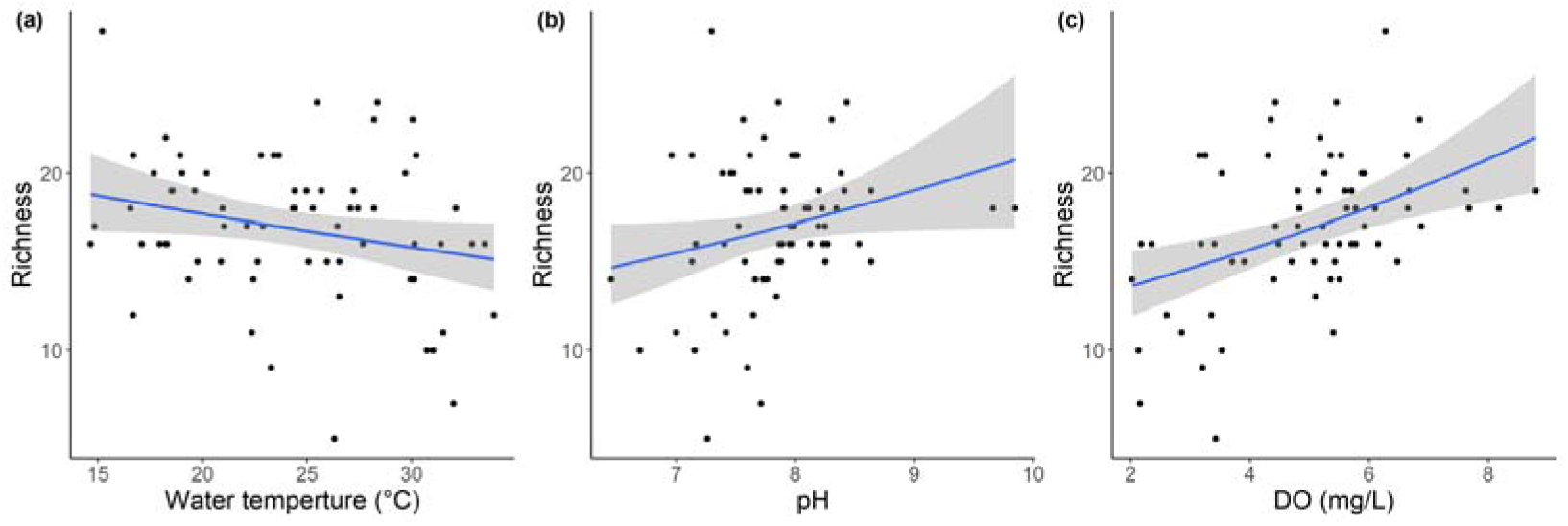
Statistically significant (*P* < 0.05) effects of environmental factors on family richness of aquatic macroinvertebrates. (a) water temperature; (b) water pH value; (c) amount of dissolved oxygen (DO).

### 3.5. Vertebrate family richness, abundance, and diversity

In total, 381 individuals of 15 vertebrate families were recorded (Appendix B). A sum of 198 individuals of 15 families were documented in organic fields, including 137 detections of 12 bird families, one detection of one reptile family, 57 detections of one amphibian family, and three individuals of one fish family. On the other hand, 183 individuals of 13 families were documented in conventional fields, including 120 detections of 10 bird families, two detections of one reptile family, 57 detections of one amphibian family, and four individuals of one fish family.

There was no significant difference in abundance, richness, and Shannon index of vertebrates between organic and conventional fields during the cultivation and fallow periods, and when both were combined (all *p* > 0.05) (Fig. 9). However, effect sizes were large for richness and Shannon index during the fallow period (1.22, 1.25, respectively). Likewise, there was no significant difference in abundance of each of the four vertebrate groups between organic and conventional fields during the cultivation and fallow periods, and when both were combined (all *p* > 0.05) (Fig. 10a-d). There was also no difference in abundance of migratory waterbirds between the practices during the fallow period (*t* = 1.52; *p* > 0.05) (Figure 10e). However, effect size of amphibian and migratory waterbirds were both large in fallow period (−1.01, 0.88, respectively).

**Fig. 9.**
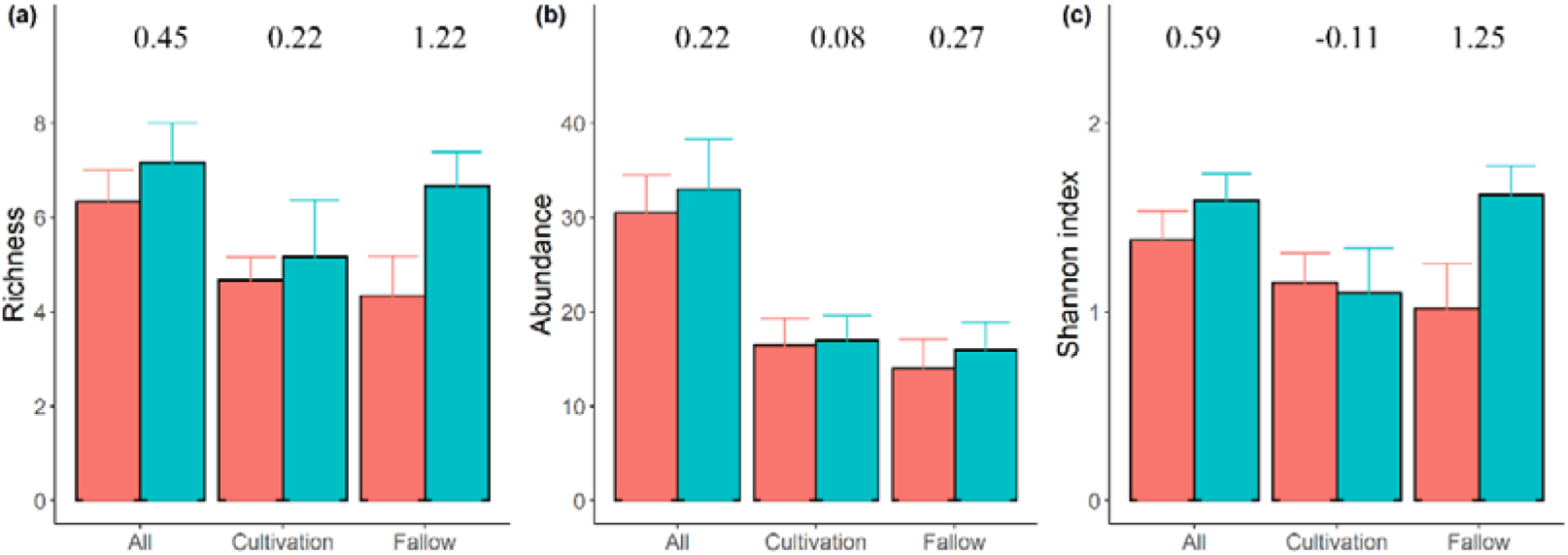
Family richness, abundance, and Shannon index of vertebrates (mean + 1 SE) in conventional (red) vs. organic (blue) rice fields during the cultivation and fallow period. (a) richness; (b) abundance; (c) Shannon index. Value above each bar represented the effect size (organic relative to conventional).

**Fig. 10.**
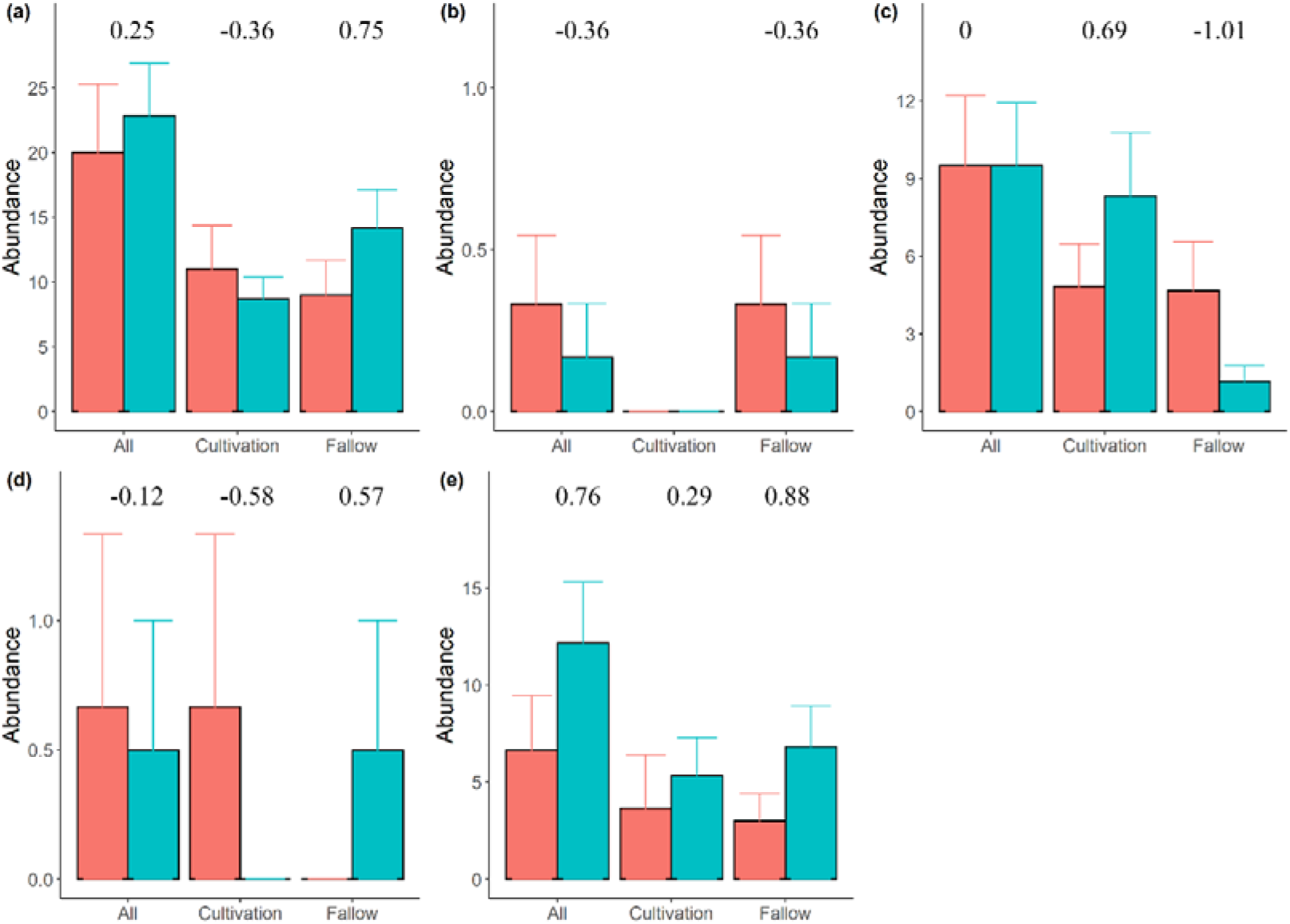
Abundance (mean + 1 SE) of five vertebrate taxa in in conventional (red) vs. organic (blue) rice fields during the cultivation and fallow period. (a) birds; (b) reptiles; (c) amphibians; (d) fish; (e) migratory birds. Value above each bar represented the effect size (organic relative to conventional).

## 4. Discussion

Overall, we observed significantly higher family richness and abundance of macroinvertebrates in organic than in conventional fields (Fig. 3). Nevertheless, the extent of difference varied with habitats (terrestrial vs. aquatic) and farming stages (cultivation vs. fallow).The differences were more remarkable for terrestrial invertebrates during the cultivation period (Fig. 3). Temporal comparisons further revealed that richness and abundance of terrestrial invertebrates increased with the progression of rice cultivation in both types of paddies, so did the differences between the two agricultural practices (Fig. 4). Richness and abundance of terrestrial invertebrates were positively associated with temperature and height of rice stands (Fig. 7), which explained why terrestrial invertebrates increased with the progression of rice cultivation from March to June (Fig. 2). Higher temperature which facilitates the growth and reproduction of invertebrates, along with taller rice stands which provide larger areas for invertebrates to inhabit or subsist on, might lead to higher number of invertebrates in later stages of cultivation period. Positive correlation of terrestrial arthropods with crop age and rice plant height has also been reported elsewhere (Bambaradeniya and Edirisinghe, 2009).

The fact that terrestrial invertebrates increased with rice cultivation stages in conventional paddies suggested that pesticides were unable to completely suppress invertebrates, including pests (Fig. 6). Because the area of rice paddies is small in Taiwan (average 1.14 ha, Huang et al. 2002), pests can easily colonize recently pesticide-applied paddies from nearby pesticide-not-yet-applied paddies. Temporal variation in pesticide applications among farmers can facilitate the maintenance of pests through metapopulation dynamics. This happens when the time window of pesticide application by different farmers is not short enough so that the pest extinction rate ls lower than the colonization rate. Coordination among farmers in delivering pesticides as synchronous as possible should therefore be helpful for controlling the pest.

On the other hand, the rate of increase in terrestrial invertebrates was higher in organic than in conventional paddies so that the difference was getting larger over the course of cultivation (Fig. 4). Such divergence might reflect the accumulated effects of pesticides on refraining the growth of invertebrate populations in conventional fields. In comparison to organic paddies where abundance of each functional group typically increased over time, number of predators and pollinators decreased during the final cultivation period in conventional paddies (Fig. 6). Predators and pollinators might therefore be more susceptible to harmful effects of pesticides, especially considering that predators usually increase following the proliferation of pests. However, it should be noted that continual increase in predators and parasitoids in organic fields was still incapable of controlling the pest population, which also reached a higher number than those in pesticide-applied conventional fields (Fig. 6). More pests can result in lower rice yield in organic than in conventional paddies. In Taiwan, yields of conventional rice per hectare averaged 7,125 and 4,417 kg (for the 1st and 2nd annual cultivation periods) while yields of organic rice averaged 4,739 (33.5% lower than conventional) and 3,345 kg (24.3% lower) (Huang et al., 2018a). Globally, yields are also generally lower for organic than conventional agriculture (Seufert et al., 2012; Smith et al., 2020). Lower organic rice productivity means more lands are needed to fulfill the same food demand as cultivated in conventional ways. A cost-effectiveness analysis of sustainability and food security should be considered in the future.

Likewise, the difference in aquatic macroinvertebrates between the two agricultural practices during the cultivation period increased from March to May (after which the lands were drained to prepare for harvest) (Fig. 4). This might similarly be due to the suppression of invertebrates by pesticides in conventional paddies. At the same time, abundance of aquatic invertebrates decreased from March to May in both types of paddies (Fig. 4). We observed lower water levels in May than in March. Less habitats might lead to fewer aquatic invertebrates in May when soils in some areas became exposed instead of flooded.

On the other hand, during the fallow period, richness of aquatic macroinvertebrate increased slowly while abundance firstly increased, followed by a small decline (Fig. 4). After the harvest, farmers introduced water to flood the fields again. This created habitats for aquatic organisms to gradually colonize to, resulting in an increase in richness over time. Therefore, negative association between aquatic invertebrate richness and water temperature (Fig. 8) might be due to seasons with higher temperature happened to be in the beginning of fallow period when richness was low (Fig. 4). Richness was positively correlated with amount of dissolved oxygen (Fig. 8), which typically decreases with temperature. Higher temperature could alternatively suppressed richness through lowering amount of dissolved oxygen. Likewise, continual colonization and lower temperature might lead to the elevation in aquatic invertebrate abundance, but it is unknown why a small decrease occurred in the end of fallow period.

As predicted, we observed larger difference in abundance between the two practices in the early stage of fallow period (effect size > 0.8), despite that difference in richness was small (effect size < 0.5) and little varied (Fig. 4). Termination of pesticide usage after rice harvest in conventional fields might partially explain the decreasing difference in abundance. Pesticide residues in the soils could gradually be diluted over the course of fallow period, with harmful effect on aquatic organisms diminishing with time. Similarly, there were more aquatic invertebrates in organic than in conventional Australian rice fields in the early part of rice season when pesticides were applied; invertebrate community became more similar with the progression of rice season (Wilson et al. 2008). In Brazil, rice paddies had fewer aquatic macroinvertebrates than natural ponds shortly after pesticide usage, but the difference diminished over time with reduced concentration of pesticide residues (Stenert et al., 2018). Due to the high costs for pesticide analysis in Taiwan, we were not allowed to assess association of temporal variation in invertebrate richness with concentrations of pesticide residues in flooded water. Given that pesticides can have disruptive effect on the whole food web (Yamamuro et al., 2019), knowledge on pesticide residues also has important implications for predators of aquatic invertebrates, including waterbirds. In northern Taiwan, fallow period (August to February) is largely overlapped with the migratory bird season (October to March), and flooded paddies provide migratory waterbirds important foraging habitats. We found no significant difference in migratory waterbird abundance between the two agricultural practices, although the effect size was large during the fallow period (> 0.8) when abundance in organic was 2.3 times that in conventional fields (mean abundance: 6.8 vs. 3 individuals; Fig. 10e). This suggests that although there was little difference in richness and abundance of aquatic invertebrates (in November and December, Fig. 4) that are potential food resources for waterbirds (Taft and Haig, 2005), other factors might lead to moderately more migratory waterbirds in organic fields. Because body size of invertebrates was not measured in this study, it is unknown whether organic fields harbored larger invertebrates. Due to that family composition of aquatic invertebrates was similar between the two practices (Fig. 5), it cannot be excluded that organic fields might shelter similar invertebrate taxa but with larger body size.

Despite that organic fields support marginally more migratory waterbirds than conventional fields, it should be stressed that these agricultural lands can be inferior habitats than natural wetlands. Lai (2012) recorded lower waterbird species richness and population density in rice paddies than in nearby wetlands in Yilan. Higher species richness and abundance of waterbirds in natural habitats than in rice fields have also been documented in other countries (Tourenq et al., 2001; Richardson and Taylor, 2003; Bellio et al., 2009). Such lower waterbird diversity in rice fields can be due to less habitat heterogeneity (Fahrig et al., 2011) or/and lower invertebrate diversity that is largely confined to resilient and opportunistic species (Bambaradeniya et al., 2004). Nonetheless, rice fields can be alternative or complementary habitats for wetland species (Elphick, 2000; Toral et al., 2012; Márquez-Ferrando et al., 2014; Pernollet et al., 2015), especially when natural wetlands have mostly been degraded.

There were more amphibians in organic fields during the cultivation period but the pattern was opposite during the fallow period (Fig. 10c). Because the Indian rice field frog (*Fejervarya limnocharis*) was the only species recorded (including tadpole and frog stages), such difference is not due to differential habitat preference of distinct amphibian species in different seasons (cultivation period in spring and summer and fallow period in fall and winter). More amphibians in conventional fields during the fallow period were chiefly due to their sheltering more frogs in November (data not shown). Lower aquatic concentration of pesticide residues in November, along with well adaptation of the Indian rice field frog to rice fields (Kuan and Lin, 2011), suggest that other factors, such as local abundance, might be more important in determining amphibian abundance that pesticide usage.

Due to limited taxonomic information on many invertebrate groups in Taiwan, organisms were identified to families instead of species. This will certainly underestimate invertebrate richness. Family richness can be a reliable surrogate to species richness (Williams and Gaston, 1994; Heino and Soininen, 2007; Slimani et al., 2019), except for sites with high species biodiversity (Balmford et al., 1996). Therefore, classification to family level should suffice given that the purpose was a comparison among agricultural practices in rice fields with generally depauperate fauna, instead of documenting the actual species richness.

In conclusion, our study provided a comprehensive picture on terrestrial and aquatic macroinvertebrate and vertebrate fauna in a representative Southeast Asia rice paddy ecosystem, at the same time considering the influence of agricultural practices (organic vs. conventional), period (cultivation vs. fallow), and environmental factors (temperature, pH value, etc.). Such a holistic approach should facilitate better evaluation of conservation value of agroecosystems and the potential risks they are exposed to, as well as help guide agricultural policy for a more sustainable solution. It is recommended that further work compares rice paddies with natural habitats (e.g. wetlands or ponds) for a finer knowledge on organisms agricultural fields can and should aim to preserve, an important prerequisite for capitalizing on agroecosystems for biodiversity conservation.

## Acknowledgements

We are indebted to laboratory colleagues for help with the field survey. This study was partially supported by Taiwan Ministry of Science and Technology (MOST 107–2311-B-003-003) awarded to CCK.

---

**Appendix A.**
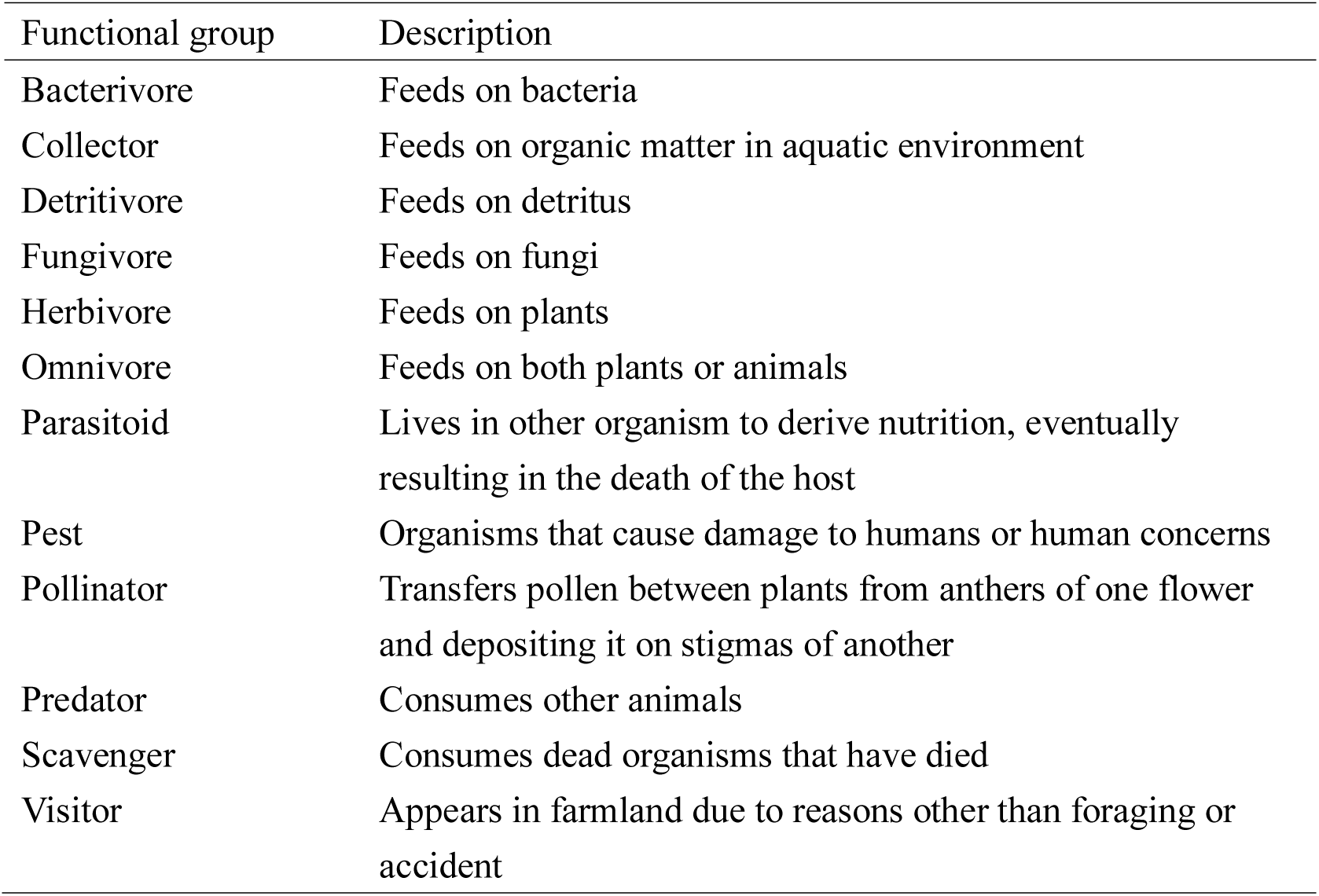
Definition of different functional groups.

**Appendix B.**
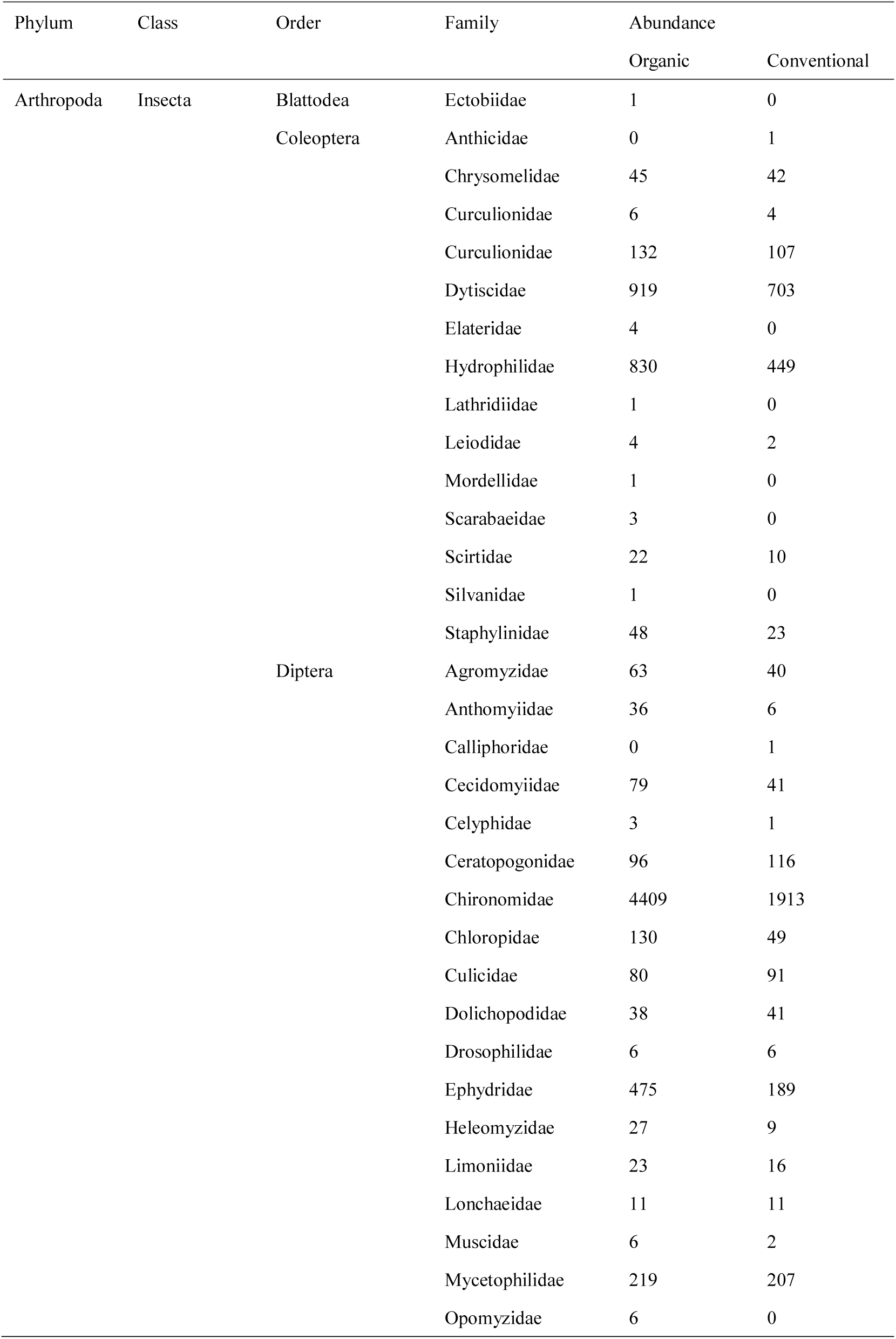
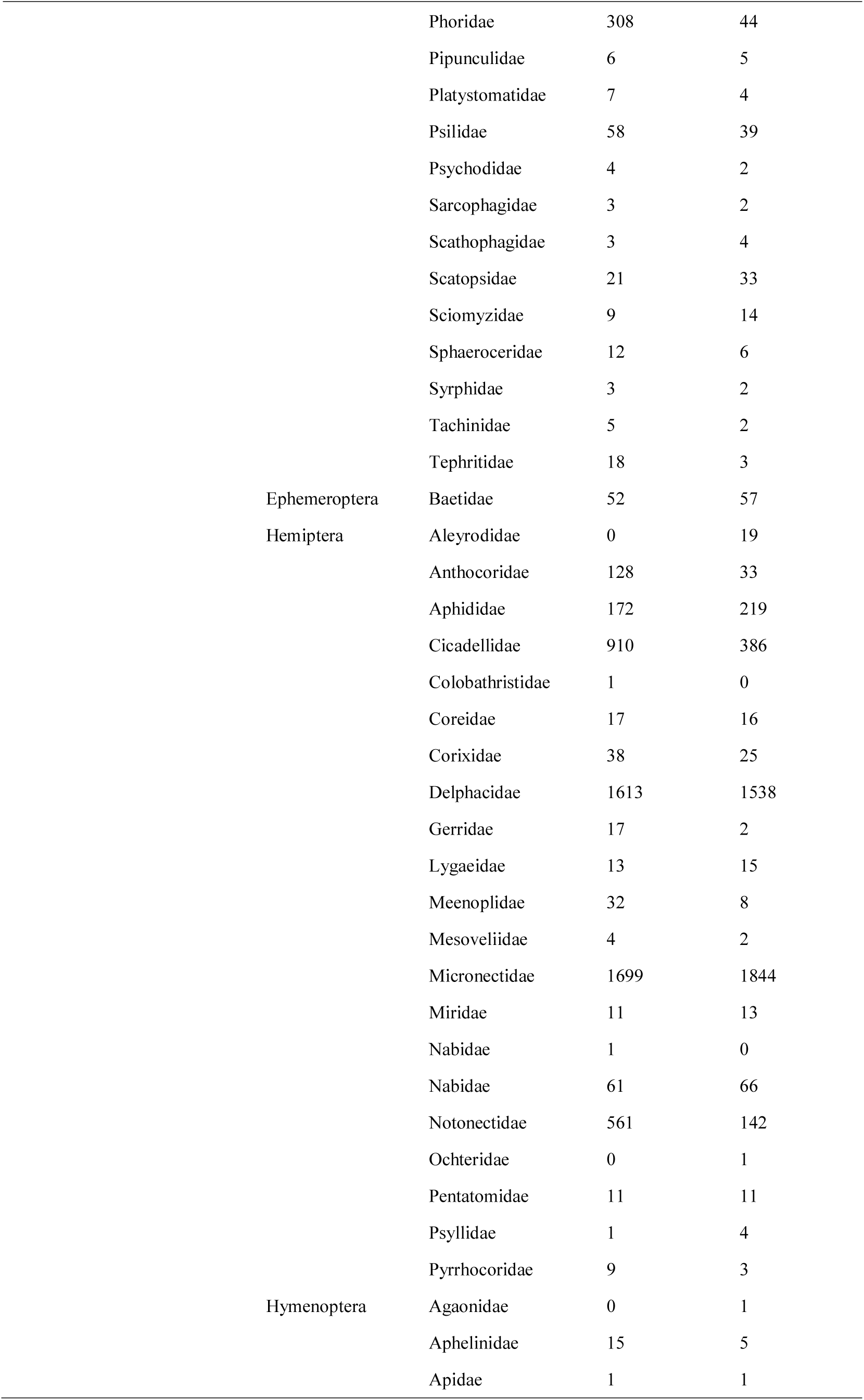
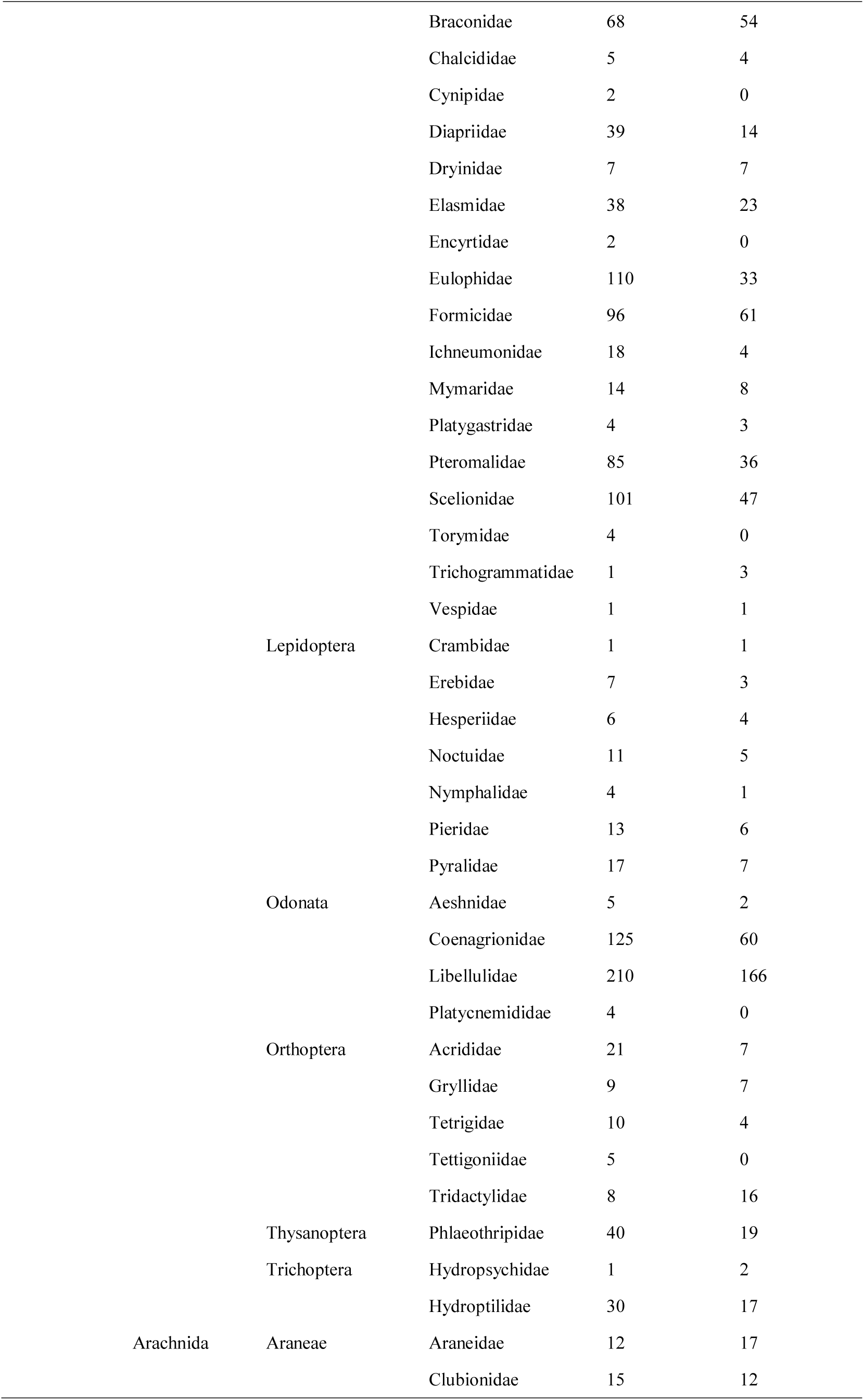
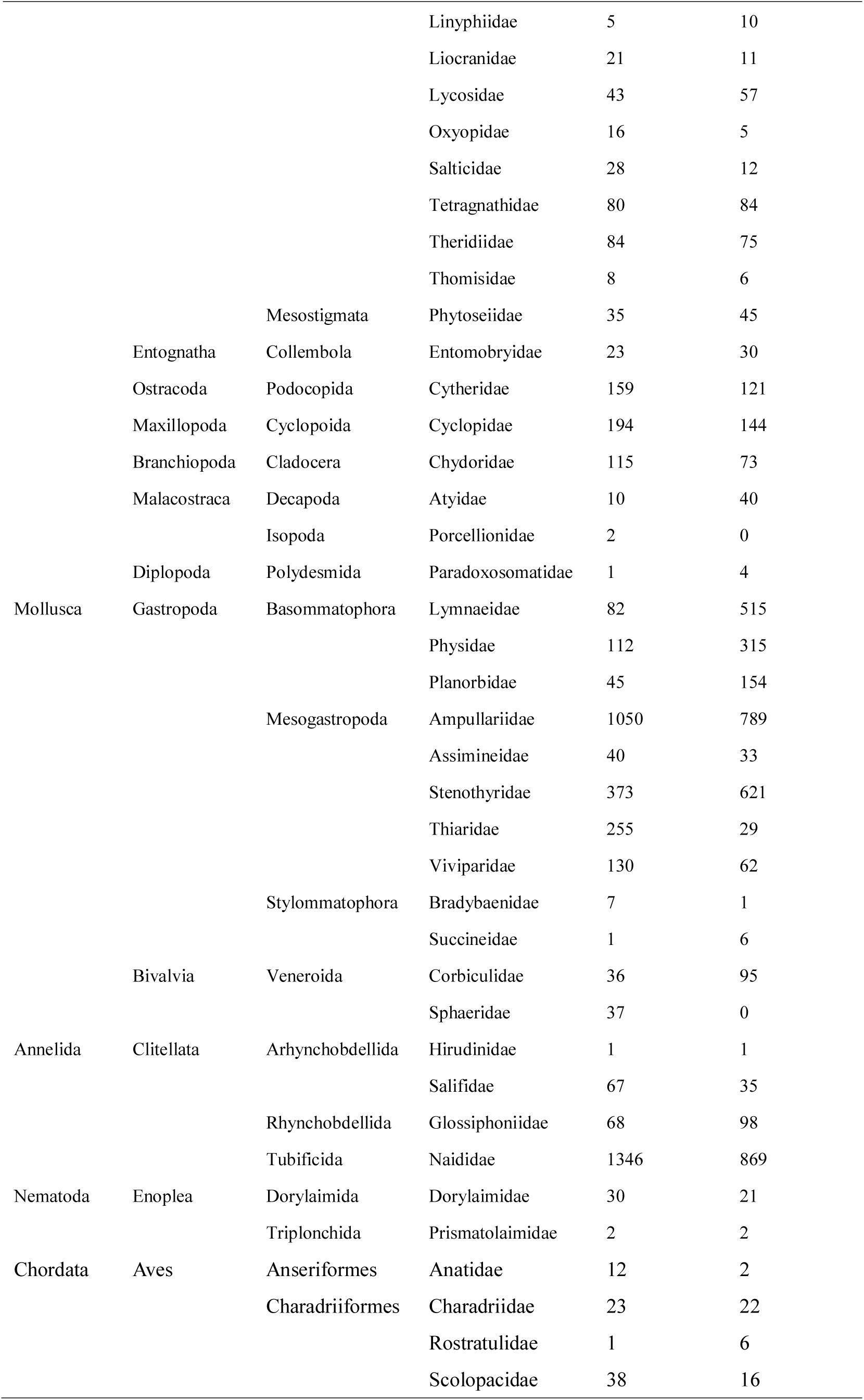
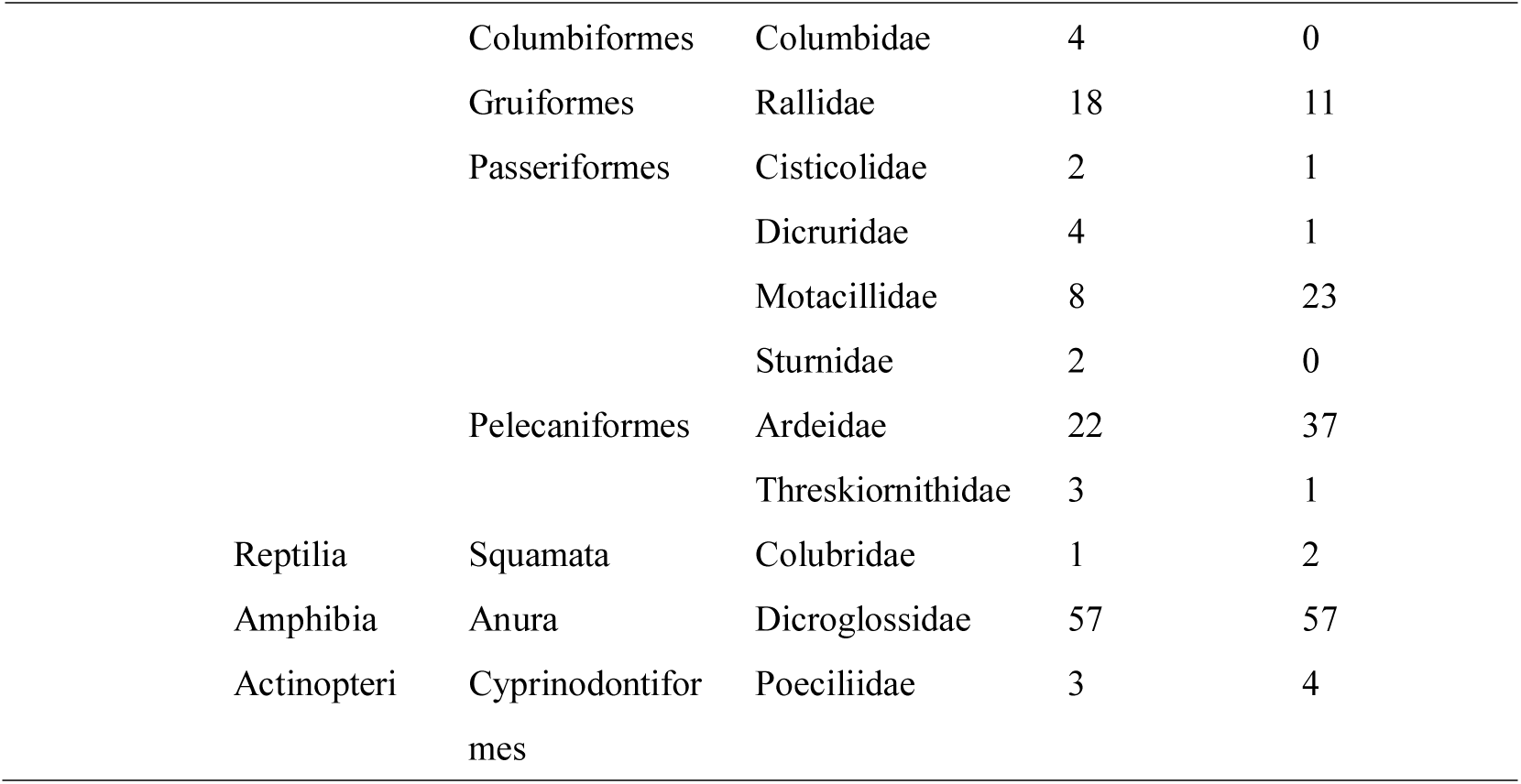
Abundance of each macroinvertebrate and vertebrate taxonomic group recorded in organic and conventional rice fields in Yilan, Taiwan.

